# REVIEW AND ANALYSIS OF MERCURY LEVELS IN BLUE MARLIN (*Makaira nigricans*, Lacepède 1802) AND SWORDFISH (*Xiphias gladius*, Linnaeus 1758)

**DOI:** 10.1101/043893

**Authors:** Tiago Rodrigues, Alberto F. Amorim

## Abstract

Mercury in the environment comes from natural processes and industrial activities. The inorganic form associates with the organic matter resulting on methylmercury (MeHg), the most toxic form to man. Mercury occurs in fishes mostly in organic form, the methylmercury. The billfishes appear to have high mercury levels because they are large predators that occupy top levels in the trophic chain. The mercury biomagnificates through the trophic chain and reaches high concentrations in top predators. The present study summarized the information about mercury levels in billfishes available in the literature, with emphasis on swordfish and blue marlin and made an analysis of data from several studies using the multivariate technique Agglomerative Hierarchical Cluster (AHC). The total mercury (THg) ranged from 0.7μg/g to 12.2μg/g in blue marlin and from 0.04μg/g to 5.1μg/g in swordfish, with data from 121 blue marlin specimens and 354 of swordfish. While a large part of the mercury content of the blue marlin was above 0.5μg/g allowed by WHO, the specimens of swordfish summarized on the present study showed generally lower values within or slightly above this limit, but the swordfish is still consumed nowadays, and large amounts consumed oftentimes may represent a risk to human health.

## 1. INTRODUCTION

### 1.1. Mercury in the environment, bioaccumulation and biomagnification

Mercury is usually found in the environment vaporized in the atmosphere, on soil, rocks, lakes, rivers, oceans and on sediment. (DIAS *et al*. 2008).

According to Kasper *et al*. (2007) some traditional uses of mercury are in the chlorine industry, manufacture of electrical devices, paints, fungicides, lamps and scientific instruments.

Contamination of the environment (atmosphere, oceans, soils and rivers) by the persistent pollutants results in a great threat to human health because these pollutants are cumulative and toxic to organisms.

Mercury is released into the environment in inorganic form and then joins the organic matter forming methylmercury (MeHg), the most toxic form to man, having a great affinity for nerve cells (Ferreira et al., 2012). According to Tavares (1995), methylmercury is the predominant form in living organisms and fish, while the inorganic mercury predominates in the atmosphere, water and soil.

Organometallic compounds of mercury have great affinity for sulfhydryl groups and hydroxyl from proteins of living beings and are very soluble in lipids, which facilitates its diffusion through the phospholipid bilayer of cell membranes, making this heavy metal easier to be absorbed and accumulated in living tissue (WHO, 1990).

In addition to accumulate in an organism (bioaccumulation), mercury also magnifies among the food chain (biomagnification) and as a result the top organisms of the chain have higher concentrations than those at the beginning of the chain (KASPER *et al*., 2007).

Due to bioaccumultaion and biomagnification, mercury levels are being transferred from one lower trophic level to a higher trophic level of the food chain, until reach the human species (CHEN, 2000; SANTOS, 2008).

### 1.2. Toxicology and effects

The mercury exposure to humans primarily occurs through food, particularly by fish consumption (SANTOS, 2008).

According to Porvari (2003) after the bioaccumulation in fish, the mercury is slowly eliminated. In addition, existing a positive relationship between fish size and mercury levels, people who consume large fish have a higher risk of mercury contamination than people who eat small fish (STORELLI, 2007). From 40% to 100% of the mercury that bioaccumulates in fish muscle tissue is methylated, making some kinds of fish not so recommended for human consumption (DIAS *et al*., 2008). Bisinoti (2004) alleges that the half-life of mercury in fish varies according to species, but usually ranges from one to three years.

The methylmercury affects the central nervous system and is a teratogenic agent, may causing genetic malformations and developmental disorders in the fetus (DIAS et al., 2008). According to Jewett (2007), pregnant women, newborns and children are the most vulnerable people to methylmercury.

Bisinoti (2004) describes the primary symptoms of neurological diseases caused by direct exposure to mercury: visual disorders such as blurred vision and reduced field of view, impaired locomotion and low motor coordination, decreased skin sensitivity, nerve pain, hearing loss, nausea and vomiting, speech problems, fatigue, weakness, and in cases of severe exposure, paralysis that can lead to death.

The investigation of mercury accumulation in fish is usually done in muscle because it is the most ingested edible tissue. On the other hand, high levels are also found in the liver, pancreas, heart and gills (SANTOS, 2008).

To evaluate if the fish is safe or not to consumption we have the limits of mercury levels stated by World Health Organization (WHO) globally and by *Agência Nacional de Vigilância Sanitária* (ANVISA) in Brazil. WHO has established the limit of 0.5*μg/g* and ANVISA stated the limit of 1.0*μg/g*.

### 1.3. Mercury in Blue marlin and Swordfish

The blue marlin (*Makaira* nigricans) belongs to the Family Istiophoridae and is one of the large billfish species, it reaches more than 800kg and 450cm (ROBINS AND RAY, 1986). In the Atlantic Ocean, common sizes are from 180cm to 300cm lower jaw to fork length (GOODYEAR AND AROCHA, 2001).

The swordfish (*Xiphias gladius*) is the only billfish that belongs to the Family Xiphiidae and is also a large fish. It can reach more than 400cm and weigh more than 500kg, although individuals weighing more than 230kg in the Mediterranean and 320kg in the Atlantic are rare (NAKAMURA, 1985).

Swordfish and blue marlin, as the other billfishes, perform extensive migration across the Atlantic being captured by several countries in different areas (Hazin et al. 1994). Additionally, the incidental capture of other billfishes by the commercial fleet added to its low market value and its wide distribution attribute difficulties for sustainable management of their stocks in the Atlantic (Uozumi and Nakano, 1994, Oliveira, et al. 2007). According to Graves and Horodysky (2010), most of the stocks of these species are over-exploited. On the other hand, billfishes are potential targets of sport fishing and have a conservational appeal, but there is not so much biological information about their stocks (ICCAT, 2005; MAYER and ANDRADE, 2009). Swordfish also has a representative capture on surface longline (DIAS *et al*., 2008).

The blue marlin and the swordfish are long-lived and large carnivores top chain fish, and their high levels of mercury due to bioaccumulation are acquired during their life from this element biomagnification throughout the trophic chain (LALLI and PARSONS, 1997).

It is well known the high levels of mercury in sharks and tunas (ADAMS *et al*., 2004). However, billfish also have great potential to accumulate this element and species such as swordfish are still consumed without any restriction, while detailed studies of this group of fish remain scarce.

The present study aimed to summarize the information about mercury levels in billfishes available in the literature, with emphasis on swordfish and blue marlin and make a single analysis of data from several studies using the multivariate technique Agglomerative Hierarchical Cluster (AHC).

## 2. MATERIAL AND METHODS

For the proposal to summarize data on mercury levels in swordfish and blue marlin were selected articles available in the literature that focused on the total mercury (THg) in the muscle tissue of these species in the Atlantic, although some additional data were also computed for enrichment of the analysis.

The literature used were Shomura and Craig (1972); Shultz and Ito (1979); Mahaffey and Rice (1998); Mendez *et al* (2001); Luckhurst *et al* (2006); Russel (2005); Dias *et al* (2008); Ferreira *et al* (2012) and Torres-Escribano *et al* (2010). All the data were analyzed in conjunction, considering the sample size N, the range and the means of the mercury levels.

The study area was also important to compare the results. As some authors have used *ppm*, *μg/g* or *mg/kg* as unit for mercury levels, for the present study *μg/g* was selected as a standard.

The multivariate technique Agglomerative Hierarchical Cluster (AHC) was performed using dissimilarities: Euclidean Distance and Ward Method with the mercury levels (minimum, maximum and means) of blue marlin and swordfish to visualize how the locations from samples were grouped. With the AHC is also possible to observe from the distance measures (Euclidean) if a group of samples from one location is closer or farther from another. All the analysis was performed using the *JMP Software* from SAS *Institute* (Version 10.0).

## 3. RESULTS AND DISCUSSION

The summarized data corresponding to the studies reviewed were basically number of individuals sampled, summary of mercury levels (minimum, maximum and mean) and the location from each group of samples.

The total number of individuals analyzed by the nine studies previously mentioned in the period between 1972 and 2012 was 475, corresponding to 121 blue marlin (*Makaira nigricans*) and 354 swordfish (*Xiphias gladius*).

The locations of origin of the fishes analyzed were Hawaii (SHOMURA and CRAIG, 1972; SHULTZ and ITO, 1979); California (SHOMURA and CRAIG, 1972); Maryland (LUCKHURST *et al*., 2006); Florida (MAHAFFEY and RICE, 1998); Bermuda (LUCKHURST *et al*., 2006); Bahamas (LUCKHURST *et al*., 2006); Gult of Mexico (LUCKHURST *et al*., 2006); Caribbean (LUCKHURST *et al*., 2006); Australia (RUSSEL, 2005); Southeast Brazil (DIAS *et al*., 2008; FERREIRA *et al*., 2012); Uruguay (MENDEZ *et al*., 2001) and Spain (TORRES-ESCRIBANO *et al*., 2010).

The range of total mercury levels in white muscle was from 0.7μg/g to 12.2μg/g in blue marlin and from 0.04μg/g to 5.1μg/g in swordfish. The highest mercury level in blue marlin (12.2μg/g) was observed by Luckhurst *et al* (2006) from Bermuda and the highest level in swordfish (5.1μg/g) was by Ferreira *et al* (2012) from Southeast Brazil.

The minimum and maximum values of mercury levels were represented in boxplots separated by species in Figure 1.

**Figure 1.**
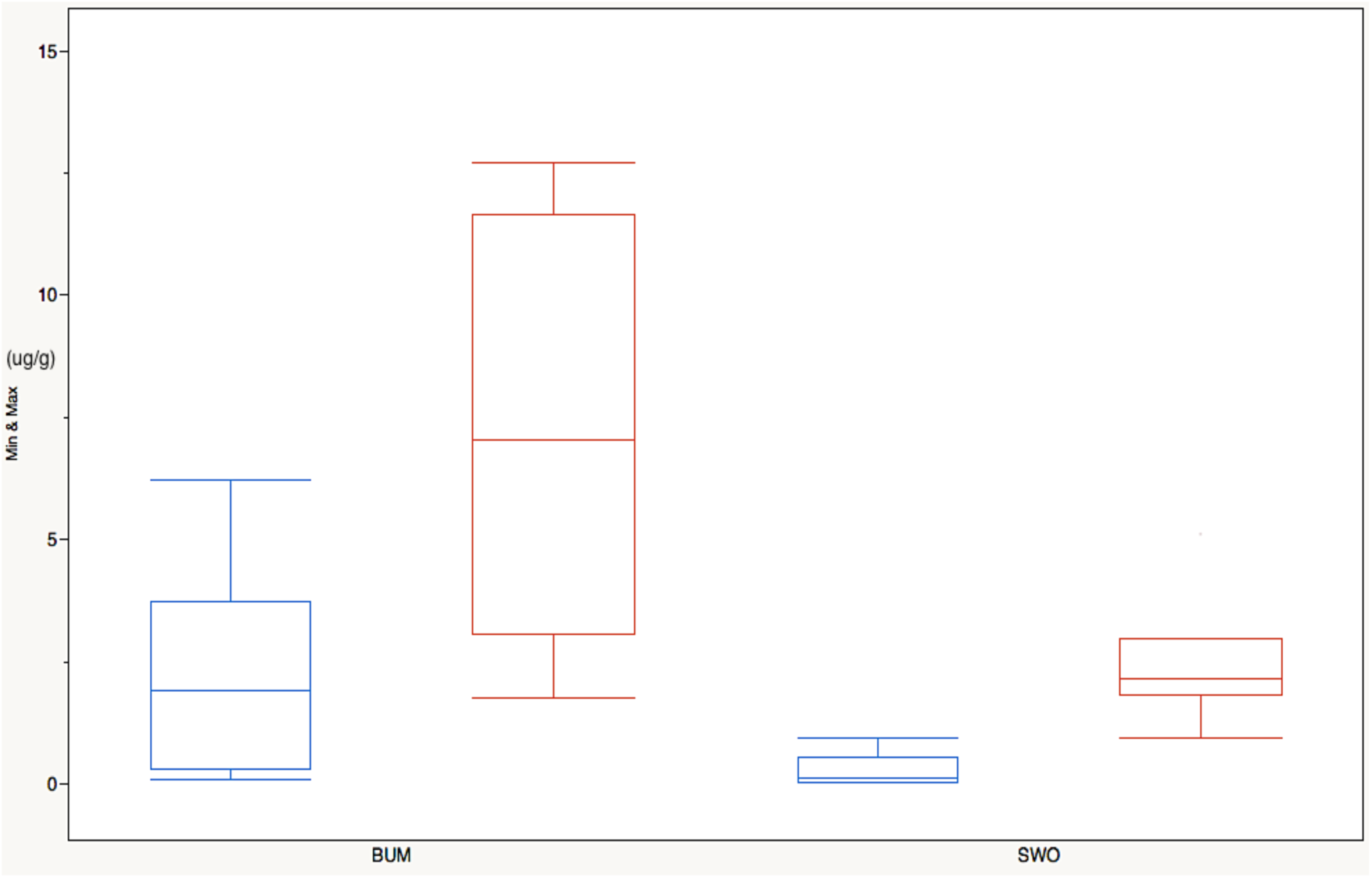
Distribution of mercury levels μg/g) in white muscle of blue marlin (BUM) and swordfish (SWO). Blue boxplot: minimum values and Red boxplot: maximum values.

Shomura and Craig (1972) also noticed high levels of mercury in blue marlin livers, with the great value of 29.55μg/g in one sample from Hawaii; probably one of the highest total mercury concentration that was ever noticed.

The means of total mercury levels in blue marlin and swordfish were grouped in Figure 2 by author, including the liver analysis of blue marlin.

**Figure 2.**
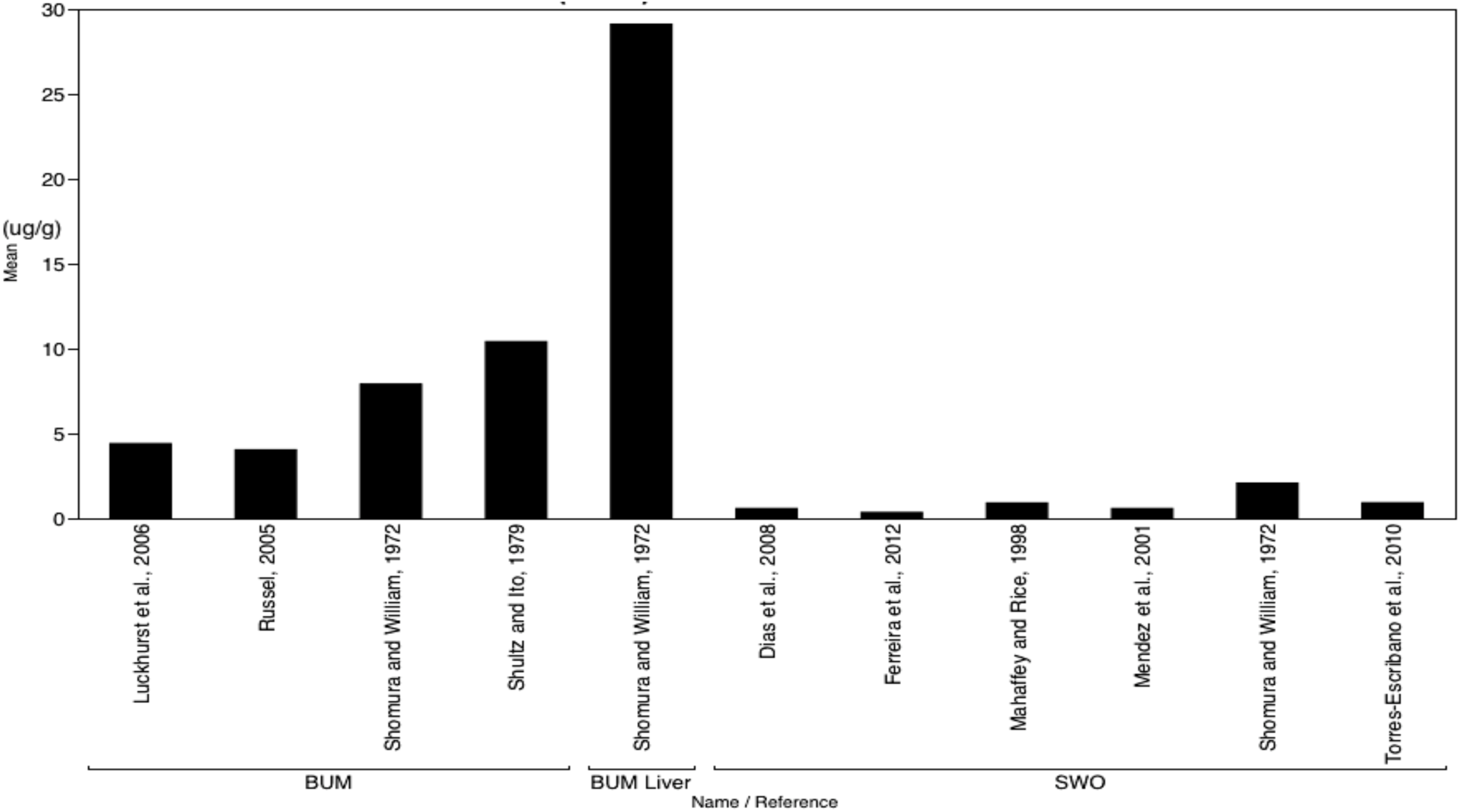
Means of total mercury levels (μg/g) in white muscle of blue marlin (BUM) and swordfish (SWO) by author. The *BUM liver* is an extra value for comparison.

The minimum values of mercury concentration in blue marlin and swordfish were from smaller individuals. As a comparison, Shomura and Craig (1972) also analyzed 14 individuals of striped marlin (*Tetrapturus audax*) from Hawaii and 42 from California; the range of total mercury was from 0.03μg/g to 2.1μg/g. The striped marlin is a smaller specie of billfish, then those lower mercury levels were expected.

The means of total mercury levels were grouped by species and location in Figure 3, again including the liver analysis of blue marlin made by Shomura and Craig (1972).

**Figure 3.**
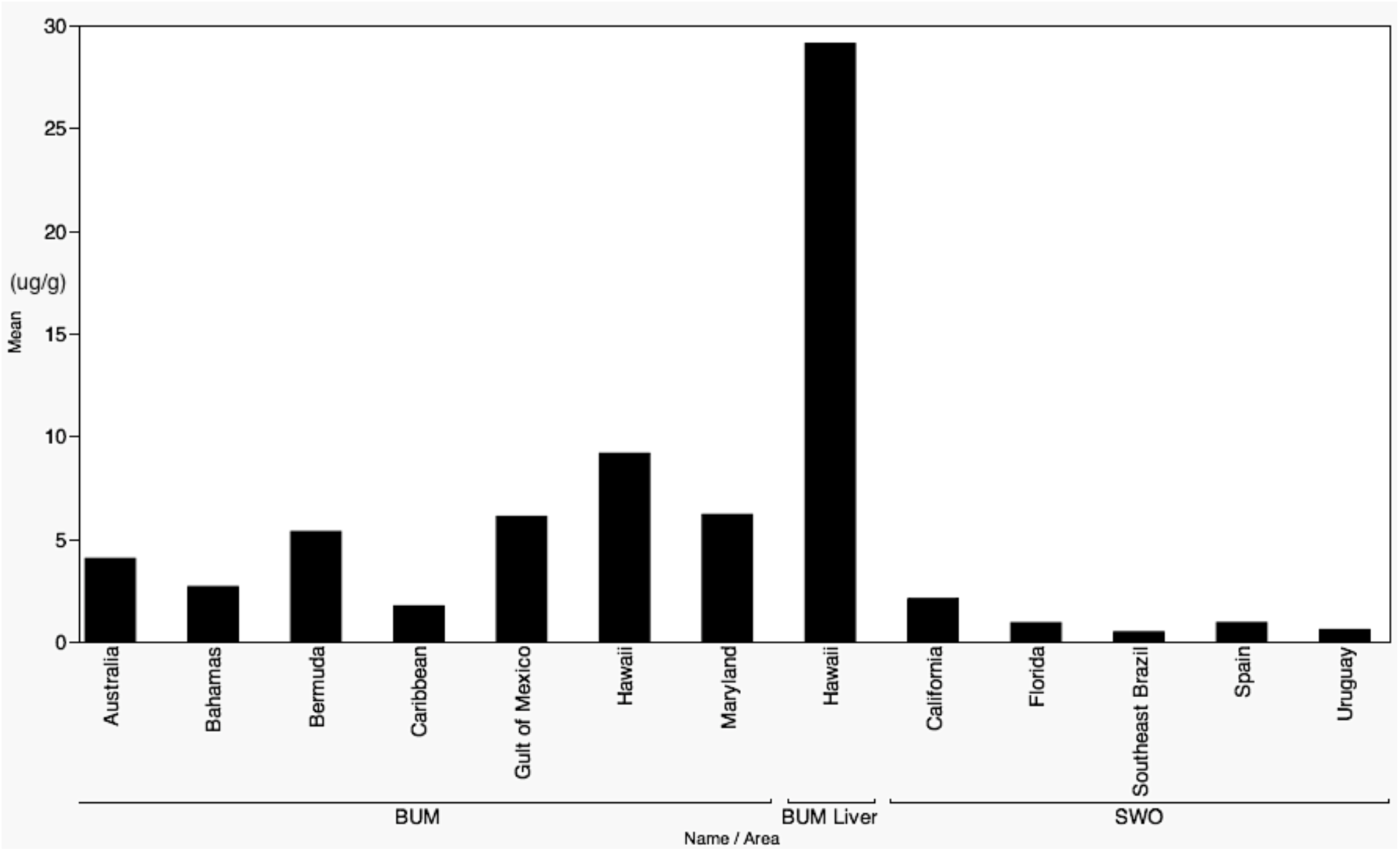
Means of total mercury levels (μg/g) in white muscle of blue marlin (BUM) and swordfish (SWO) by location. The *BUM liver* is an extra value for comparison.

The Agglomerative Hierarchical Cluster (AHC) resulted in three main groups: the first one containing both sample collection from Hawaii, Bermuda and Gult of Mexico; the second group with California, Southeast Brazil, Uruguay, Spain, Caribbean and Florida and the third group with samples from Maryland, Bahamas and Australia. The Figure 4 shows the AHC dendogram and a small distance map, so it is possible to notice the dissimilarity level, wich is higher on the right side of the figure and lower on the left side (nearby groups). The associations within the groups (noticeable in the dendogram by the linkages) represent that their values of total mercury in blue marlin and swordfish were more similar than in other location where the linkage is not so close.

**Figure 4.**
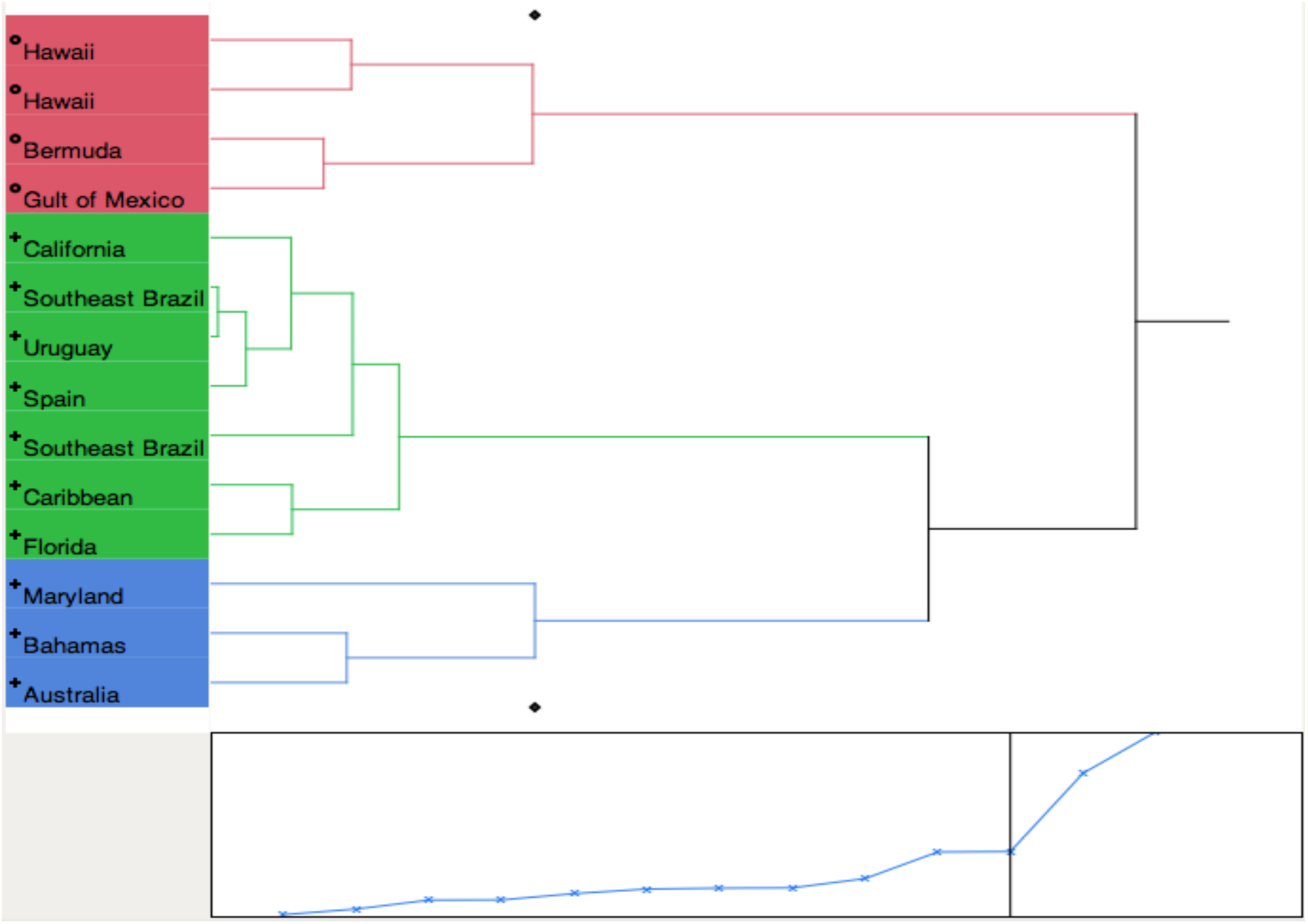
Agglomerative Hierarchical Cluster (AHC) dendogram (top) and distance map.

The first group was farther from the others, maybe because this group included samples from Hawaii (not in Atlantic Ocean like major of samples) and Bermuda and Gult of Mexico also represented high levels (see Figure 2).

Analyzing the correlation between the total mercury level and weight or length, Dias *et al* (2008) was the only study where the relationship was negative. According to them, this was probably due to the small range of lengths analyzed (DIAS *et al*., 2008). In all other studies the relationship was positive. Shultz and Ito (1979) also found a significant difference between males and females.

While a large part of the mercury content of the blue marlin was above 0.5μg/g allowed by WHO, the specimens of swordfish summarized on the present study showed generally lower values within or slightly above this limit. Dias *et al* (2008) found that 54% of all samples of swordfish were above the WHO limit. It is also important to say that the swordfish is still widely consumed currently, and large amounts consumed oftentimes may represent a risk to human health. Morgan *et al* (1997) further states that mercury is not removed from fish tissues by any practical cooking method.

Hightower and Moore (2003) studied the mercury levels in peripheral blood and hair from 123 patients and correlated this levels to fish consumption. The highest five levels reported in this study were made in five different labs and all five patients had consumed swordfish (HIGHTOWER and MOORE, 2003).

Ferreira *et al* (2012) argued that health authorities in Brazil should alert frequent consumers of swordfish due to high mercury levels in this species. This is a challenge that needs to happen worldwide, primarily in high-risk communities that consume large fishes frequently, such as tunas, billfishes and sharks.

## 4. CONCLUSION

It was concluded that despite the lower levels of total mercury in swordfish comparing it to the blue marlin, consumers need to be better informed about the risks of eating this fish frequently, as well as other species that bioaccumulates mercury, specially pregnant women.

It was also noted the lack of information about mercury levels in billfishes and further studies would be highly recommended.

## REFERENCES

Adams, D. H. 2004. Total mercury levels in tunas from offshore waters of the Florida Atlantic coast. Mar. Pollut. Bull. 49: 659–667.

Bisinoti, M. C., Jardim, W. F.; 2004. O Comportamento do Metilmercúrio (MetilHg) no Ambiente. Quim. Nova 27(4): 593-600.

Chen, C. Y., Stemberger, R.S., Klaue, B., Blum, J.D., Pickhardt, P.C. and Folt, C.L. 2000. Limnology and Oceanography 45(7): 1525-1536.

Dias. A.C.L; Guimarães, J.R.D.; Malm, O. and Costa, P.A.S. 2008. Mercúrio total em músculo de cação Prionace glauca (Linnaeus, 1758) e de espadarte Xiphias gladius (Linnaeus, 1758), na costa sul-sudeste do Brasil e suas implicações para a saúde pública. Cad. Saúde Pública, v. 24, n. 9, p. 2063-2070.

Ferreira, M. S.; Mársico, E. T.; Mársico, E. T.; Marques-Junior, A.N.; Mano, S.B.; São Clemente, S. C. and Conte-Junior, C.A. 2012. Mercùrio total em pescado marinho no Brasil. R. bras. Ci. Vet., v. 19, n. 1, p. 50-58.

Graves, J. E., and Horodysky, A.Z. 2010. Asymmetric conservation benefits of circle hooks in multispecies billfish recreational fisheries: a synthesis of hook performance and analysis of blue marlin post-release survival. Fishery Bulletin. 108:433-441.

Hazin, F. H. V., R. Lessa, R. R. Arraes, M. R. Coimbra, R. C. Souza, M. Natalino and P. S. Pantoja. 1994. Distribution and relative abundance of tunas and billfishes in the southwestern equatorial Atlantic. Inter. Comm. Conser. Atl. Tunas, Coll. Vol. Sci. Pap., 41:309-324.

Hightower, J.M. and Moore, D. 2003. Mercury Levels in High-End Consumers of Fish. Environmental Health Perspectives. 111(4): 604-608.

ICCAT (International Commission for the Conservation of Atlantic Tunas). 2005. Report of the Standing Committee on Research and Statistics. Int. Comm. Cons. Atl. Tunas, Madrid, Spain, 224 p.

Jewett, S. C., Duffy, L. K. 2007. Mercury in fishes of Alaska, with emphasis on subsistence species. Science of the Total Environment 387: 3-27.

Kasper, D.; Borato, D.; Palermo, A.F.A. and Malm, O. 2007. Mercúrio em peixes: fontes e contaminação. Oecol. Bras. V.11, n. 2, p. 228-239.

Lalli, C.M. and Parsons, T.R. 1997. Biological oceanography, an introduction. 2nd Ed. Milton Keynes: The Open University.

Luckhurst, B.E., E. D. Prince, J. K. Llopiz, D. Snodgrass, and E. B. Brothers. 2006. Evidence of blue marlin (Makaira nigricans, Lacepede, 1803) spawning in Bermuda waters and elevated mercury levels in large specimens. Bulletin of Marine Science. 79(3): 691-704.

Mahaffey, K.R. and Rice, G.E. 1998. Environmental Protection Agency Office of Air Quality Planning and Standards. Mercury Study Report to Congress. Govt Reports Announcements and Index (GRA and I), Issue 09, Available: http://www.epa.gov.

Mayer, F.P. and Andrade, H.A. 2009. Estimativa de abundância do agulhão-branco (Tetrapturus albidus) capturado no Atlêntico Sul através de modelos de contagem considerando superdispersão e excesso de zeros. Anais do III Congresso Latino Americano de Ecologia. São Lourenço.

Mendez, E.; Giudice, H.; Pereira, A.; Inocente, G. and Medina, D. 2001. Total mercury content: fish weight relationship in swordfish (Xiphias gladius) caught in the southwest Atlantic Ocean. J Food Compost Anal. 14:453-60.

Morgan, J.N.; Berry, M.R. and Graves, R.L. 1997. Effects of commonly used cooking practices on total mercury concentration in fish and their impact on exposure assessments. J Expo Anal Environ Epidemiol 7(1):1, 19–33.

Nakamura, I. 1985. FAO species catalogue. Vol. 5. Billfishes of the world. An annotated and illustrated catalogue of marlins, sailfishes, spearfishes and swordfishes known to date. FAO Fish. Synop. 125(5):65 p.

Oliveira, I.M., Hazin, F.H.V.; Travassos, P.; Pinheiro, P.B. and Hazin, H.V. 2007. Preliminary results on the reproductive biology of the white marlin, Tetrapturus albidus Poey 1960, in the western equatorial Atlantic Ocean. ICCAT Collect. Vol. Sci. Pap. 60(5): 1738-1745.

Porvari, P. 2003. Sources and fate of mercury in aquatic ecosystems. Monographs of the Boreal Environment Research 23: 1-52.

Robins, C.R. and Ray, C.G. 1986. A Field Guide to Atlantic Coast Fishes of North America. Houghton Mifflin: Boston.

Russell, J. 2005. Mercury levels in three species of marlin from eastern Australian. 4th Int. Billfish Symp., Catalina Island.

Santos, H. I. O. 2008. Acumulação de mercúrio em diferentes tecidos de cacao. Dissertation. Aveiro, Portugal.

Shomura, R.S. and Craig, W.L. 1972 Mercury in several species of billfishes taken off Hawaii and Southern California. NOAA Technical Report. SSRF-675.

Shultz, C.D. and Ito, B.M. 1979. Mercury and selenium in blue marlin, Makaira nigricans from the Hawaiian Islands. Fishery Bulletin. 76(4): 872-879.

Storelli, M. M., Barone, G., Piscitelli, G., Marcotrigiano, G.O. 2007. Mercury in fish: Concentration vs. fish size and estimates of mercury intake. Food Additives and Contaminants:1-5.

Tavares, C. M. O. F. 1995. Contaminação por Hg do Solo e Plantas dos Campos Marginais do Esteiro de Estarreja, Universidade de Aveiro. Dissertation.

Torres-Escribano, S.; Vélez, D. and Montoro, R. 2010. Mercury and methylmercury bioaccessibility in swordfish. Food Additives & Contaminants. 27(3): 327-337.

Uozumi, Y. and Nakano, H. 1994. A historical review of Japanese longline fishery and billfish catches in the Atlantic Ocean. ICCAT Coll. Vol. Sci. Pap. Madrid, 41:233-243.

WHO: World Health Organization. 1990. Methylmercury. Environmental Health Criteria, Genova.

